# Systematic review and meta-analysis of the prevalence of Strep A *emm* clusters in Africa to inform vaccine development

**DOI:** 10.1101/2020.05.06.081927

**Authors:** Taariq Salie, Kelin Engel, Annesinah Moloi, Babu Muhamed, James B Dale, Mark E Engel

## Abstract

**Background:** An *emm*-cluster based system was proposed as a standard typing scheme to facilitate and enhance future studies of Group A Streptococcus (Strep A) epidemiological surveillance, M protein function and vaccine development strategies. We provide an evidence-based distribution of Strep A *emm* clusters in Africa and assess the potential coverage of the new 30-valent vaccine in terms of an emm cluster-based approach.

**Method:** Two reviewers independently assessed studies retrieved from a comprehensive search and extracted relevant data. Meta-analyses were performed (random effects model) to aggregate *emm* cluster prevalence estimates.

**Results:** Eight studies (n=1,595 isolates) revealed the predominant *emm* clusters as E6 (18%, 95% confidence interval (CI), 12.6; 24.0%), followed by E3 (14%, 95%CI, 11.2; 17.4%) and E4 (13%, 95%CI, 9.5; 16.0%). There is negligible variation in *emm* clusters as regards regions, age and socio-economic status across the continent. Considering an *emm* cluster-based vaccine strategy, which assumes cross-protection within clusters, the 30-valent vaccine currently in clinical development, would provide hypothetical coverage to 80.3% of isolates in Africa.

**Conclusion:** This systematic review indicates the most predominant Strep A *emm* cluster in Africa is E6 followed by E3, E4 and D4. The current 30-valent vaccine would provide considerable coverage across the diversity of *emm* cluster types in Africa. Future efforts could be directed toward estimating the overall potential coverage of the new 30-valent vaccine based on cross-opsonization studies with representative panels of Strep A isolates from populations at highest risk for Strep A diseases.

**Importance:** Low vaccine coverage is of grave public health concern, particularly in developing countries where epidemiological data are often absent. To inform vaccine development for group A streptococcus (Strep A), we report on the epidemiology of the M Protein emm clusters from Strep A infections in Africa, where Strep A-related illnesses and their sequalae including rheumatic fever and rheumatic heart disease, are of a high burden. This first report of emm clusters across the continent indicate a high probably of coverage by the M Protein-based vaccine currently undergoing testing, were an emm-cluster based approach to be used.

## Introduction

Group A Streptococcus (Strep A) causes a range of human infections including pharyngitis and impetigo, which can lead to non-suppurative (immune-mediated) sequelae such as acute rheumatic fever (ARF) and rheumatic heart disease (RHD) if not properly managed (2). Additionally, Strep A has the ability to cause invasive infection such as sepsis, necrotizing fasciitis, pneumonia, and streptococcal toxic shock syndrome (STSS) in children and adults (3) with a high fatality rate; furthermore, it is a leading cause of maternal death in some regions (4). Strep A infections mostly affect young children and women living in developing countries (5). The estimated symptomatic Strep A pharyngitis annual incidence rate is 0.4 cases per person-year, with over 423 million cases, in children residing in developing countries (2).

The dire complications and huge economic burden of Strep A infections support the urgent need for an effective vaccine that would provide broad coverage of circulating Strep A strains (6). One of the Strep A vaccine strategies targets the M-protein on the bacterial surface, which has thermal stability, anti-phagocytic properties and the capacity to evoke antibodies with the greatest bactericidal activity (7). The hypervariable N-terminal region of the M-protein displays extensive nucleotide differences, thus giving rise to various M-protein amino acid sequences which imparts serological specificity (8). The 5’ *emm* sequence encoding the mature protein is the basis for categorizing different Strep A strains through molecular typing methods, which aid in defining the epidemiology of Strep A infections.

A 30-valent N-terminal M protein-based vaccine (9) is undergoing clinical trials (10). The vaccine composition was based on extensive Strep A surveillance data from developed regions such as USA and Europe, those isolates that are involved in invasive disease, those associated with superficial infections and those causing autoimmune diseases (11, 12). However, given the >200 Strep A *emm* types characterized to date (13), it is not surprising that there are highly prevalent Strep A subtypes absent in the current vaccine formulation, thus possibly excluding at-risk populations outside of western countries (14).

An *emm* clustering system was introduced by Sanderson-Smith and colleagues that phylogenetically analyzed the whole M protein sequences, organizing *emm* types into clusters that have the same or similar sequences and host protein binding properties (15). This proposed classification allows for the previously identified Strep A *emm-* types to be categorized into 48 discrete *emm* clusters (15) where more than one *emm* type may be contained within a cluster (Table 1). The *emm* cluster system compliments the *emm* typing system, which may serve to enhance studies relating to M protein function, streptococcal virulence, epidemiological surveillance, and vaccine development (15). *Emm* clusters E1-E6 were placed into clade X, binding to immunoglobulin and C4BP. While A-C1 through A-C5 and D1-D5 were grouped into clade Y, with a host protein tropism towards plasminogen and fibrinogen.

**Table 1.**
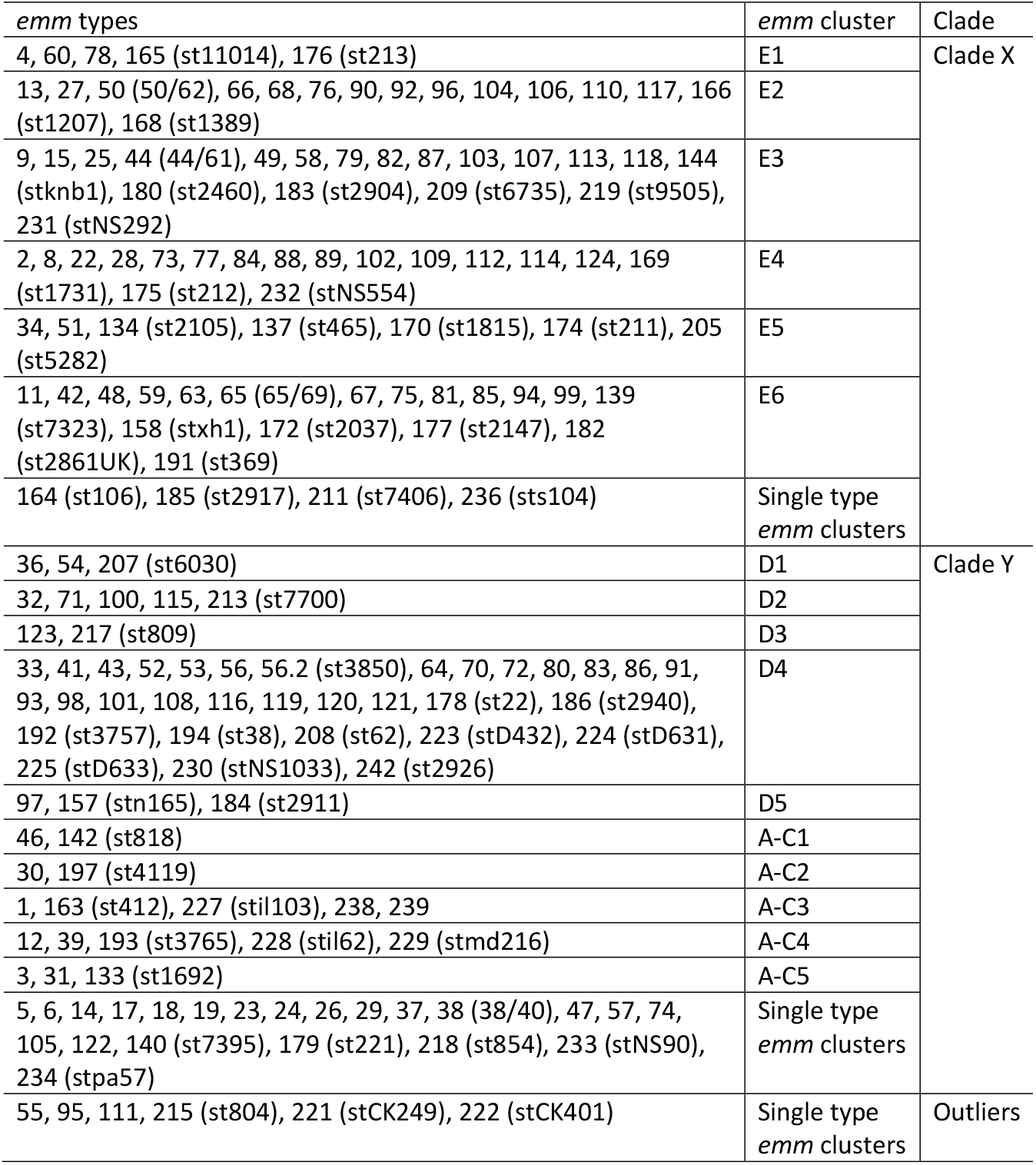
The *emm* clusters and their corresponding *emm* types. Adopted from Sanderson-Smith (2014)

To date, significant *emm* cluster data have been produced through *emm* typing of Strep A, with recent studies reporting on *emm* cluster epidemiology. Shulman documented the most prevalent *emm* clusters in the USA as E4 (27.16%), A-C3 (17.78%) and A-C4 (17.56%) amongst 7,040 isolates (16). The prevalence of *emm* clusters in three Pacific countries, viz. Australia, Fiji and New Caledonia illustrated that 70%-84% of clusters from isolates were shared, as opposed to comparison of *emm* types having only 14%-30% commonality between countries (17). In a third study by Chang-Ni in Taiwan, an analysis of both invasive and non-invasive strains revealed that cluster E6 was associated with both types of infections, while clusters D4, E2 and E3 were responsible for causing invasive isolates in their population(18). Recently, Frost demonstrated that M type–specific and cross-reactive immune responses frequently align with *emm* clusters, raising new opportunities to design multivalent vaccines with broad coverage (19).

A thorough review of *emm* cluster data from Africa has not yet been undertaken. A study that aggregates the African data on clusters is essential to contribute to the growing literature in efforts to develop a Strep A vaccine on a global scale, particularly in low-income countries where the burden of disease is greatest. Therefore, this review sought to provide an evidence-based distribution of Strep A *emm* clusters in Africa.

## Methods

This study employed rigorous methods drawn from the scientific techniques and guidelines offered by the Cochrane Collaboration (20) and by reviews published previously (21, 22). The review protocol has been registered in the PROSPERO International Prospective Register of Systematic Reviews CRD42017062485.

### Review Question

This review asks the following question: What is the prevalence of Strep A *emm* clusters in Africa in the current available literature? Is there variation in *emm* cluster prevalence based on geography, age, clinical manifestation or socio-economic status? We further sought to explore the potential coverage of the current 30-valent vaccine using a cluster-based approach.

### Search Strategy

A comprehensive strategy was developed to search electronic databases to maximize sensitivity (Table S1-Appendix). The search strategies incorporated both free term text that are controlled to suit specific databases individually and Medical Subject Headings (MeSH) adapted to suit each individual database. A combination of terms relating to *“emm* typing”, *“emm* clusters”, “*emm*/M protein” and “streptococcal diseases” focusing on the African continent by applying the African search filter previously used by Pienaar and colleagues (23). The following databases were searched as at 29 April 2020; PubMed, Scopus and Google Scholar for grey literature. The search was not restricted to any publication dates or language (however, abstracts must be clearly written in English for the study to be considered). Published and unpublished data were also considered for inclusion.

### Inclusion criteria

All studies that described the prevalence of *emm* clusters or *emm* types within a given population were included in the review. Participants were restricted to the African continent but were not discriminated by clinical manifestation of Strep A or site of Strep A isolation. All laboratory-confirmed Strep A isolates were molecularly characterized by the *emm* typing method to ascertain serotypes as this is the gold standard technique (24). The *emm* typing method as developed by Beall (25) and in alignment with the Centers for Disease Control and Prevention (26), is able to classify Strep A serotypes which is based on sequence analysis of PCR products of the 5’ hypervariable region of the M protein gene.

Two reviewers applied the search strategy to the relevant databases independently in which the titles and abstracts were evaluated to exclude studies that did not describe the prevalence of Strep A. Thereafter, full texts of the included titles and abstracts were retrieved and further evaluated against the inclusion criterion (Table S2-Appendix). A comparison was made between individual lists, if the reviewer’s lists were not concurrent, discrepancies were discussed and an arbitrator (third reviewer) was contacted to resolve any disagreements.

### Exclusion criteria

Case reports, narrative reviews, opinion pieces and publications lacking prevalence primary data, or referenced methodology according to Beall (25), were excluded from the review. Duplicated studies of the same datasets and participants were removed and the final most recent publication of the data was considered for inclusion.

### Data extraction and management

Two reviewers extracted data using a standardized data extraction form and any contradictions were solved through discussion or that of a third reviewer. Search results from the databases listed above, published and unpublished studies were managed with Endnote X9 referencing software. Briefly, data extraction consisted of recording the study demographics (amount of study participants, the geographical region, age group of enrolled participants, the clinical manifestation of disease and socio-economic status) along with the relevant *emm* type/cluster distributions within the population. Socio-economic status for the study settings was determined at a country level, according to The World Bank (27).

### Quality assessment

The risk of bias assessment established by Hoy (28) and modified by Werfalli (22), was adapted in questions specific for use in this review (Table S3-Appendix). Using a quantitative scoring system, studies were characterized being of a low, moderate or high risk of bias. A study with low risk of bias is of high-quality and a low-quality study is associated with a higher risk of bias. Assessing the risk of bias informs the evaluation of heterogeneity in the pooled analyses.

### Analysis

Data synthesis included three steps: (1) characterizing the study demographics (2) documenting *emm* types for *emm* cluster calculations, and (3) assessing potential vaccine coverage. In each study, the prevalence of *emm* types was recalculated by analyzing figures and tables to confirm the authors results and findings and to document the numerators and denominators. In older studies, *emm* typing information needed to be updated using the CDC database (29). Where *emm* cluster information was not reported, the CDC classification system was used to augment missing data (https://www.cdc.gov/groupastrep/lab.html), as well as the original cluster descriptions (15).

To calculate potential coverage, three tiers were assessed: 1) M peptides in the vaccine, 2) *emm* types that have been shown to be cross-opsonized, and 3) *emm* types that just happen to be in a cluster that are represented by one or more vaccine *emm* types. Quantitative data analysis was completed using Stata version 14.1 (StataCorp, College Station, TX, USA). We applied the Freeman-Tukey double arcsine transformation option using the *metaprop* routine to describe the combined prevalence estimates of all included studies with the standard error across the unadjusted estimates (30). *Emm* cluster distribution was correlated against different variables (resource setting, clinical manifestation and age group) in each of the studies. Lastly, we determined the theoretical protective coverage by *emm* cluster cross-opsonization for *emm* types included in the M protein-based vaccine (11).

## Results

The literature search for articles was reported according to the Preferred Reporting Items for Systematic reviews and Meta-Analysis (PRISMA) Statement (1). Figure 1 details the search results with the retrieval of 121 articles for consideration from the respective electronic databases. After title screening and the removal of duplicates, we excluded 23 articles. We reviewed the remaining abstracts and excluded a further 81 articles, leaving 17 articles requiring full-text evaluation. Finally, eight articles met the inclusion criteria and were included in the review. A list of the excluded studies with reasons are detailed in Table S4 (Appendix).

**Figure 1.**
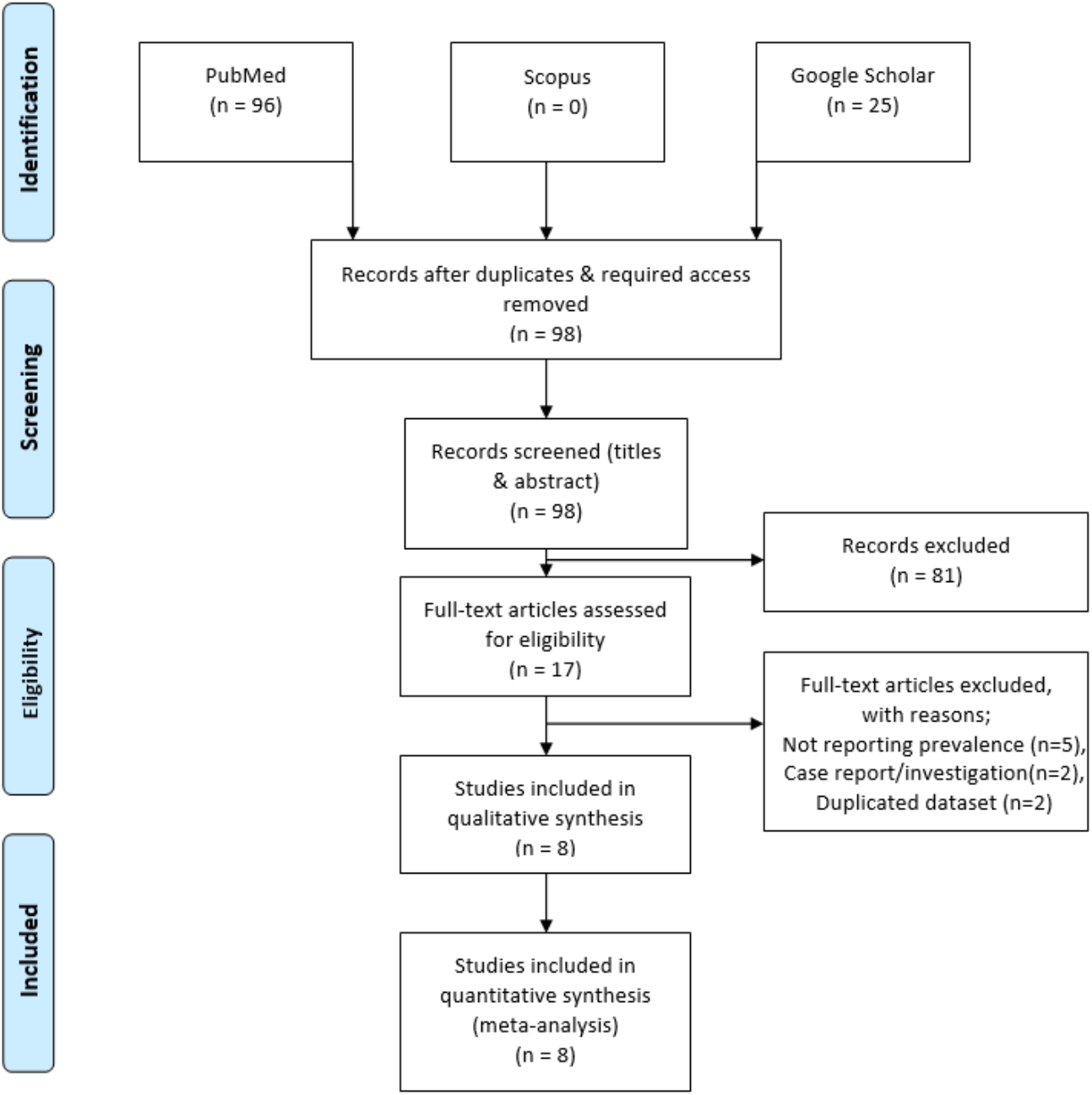
Schematic PRISMA flow diagram of the literature search (1).

### Characteristics of included studies

The included articles were published between 2004 and 2019 with sample sizes ranging from 43 and 396 total isolates. Of these, two articles had cross-sectional study designs, while the remaining studies took a prospective passive surveillance approach. The ages of participants included in the studies were also recorded; six articles studied isolates obtained from children (range 0-18 years old) and two, studied patients of all ages. Studies were conducted in local and university hospitals, clinics, outpatient departments and schools situated in the study areas (Table 2).

**Table 2.** Characteristics of included studies

| Study ID | Country | Design | Setting (Local, social context) | Socio-economic setting | Population description | Inclusion criteria | Age (years) |
| --- | --- | --- | --- | --- | --- | --- | --- |
| Abdissa 2005 | Ethiopia | Cross-sectional | Public primary schools situated in Addis-Ababa, Gondar & Dire-Dawa | Low income | Healthy children attending public primary schools | Healthy school children in area | 4-16 |
| Barth 2019 | South Africa | Prospective passive surveillance | Groote Schuur Hospital in Cape Town | Upper middle income | Children attending public clinics | Patients with confirmed Strep A infection | ALL |
| Engel 2014 | South Africa | Cross-sectional | Langa Clinic, Vanguard Community Health Centre | Upper middle income | Children attending public clinics | Children with sore throat | 3-15 |
| Hraoui 2011 | Tunisia | Passive surveillance | Microbiology Lab of Charles Nicolle University Hospital in Tunis | Lower middle income | Patients attending the local hospital | Patients with confirmed Strep A infection | ALL |
| Mzoughi 2004 | Tunisia | Prospective surveillance | Farhat Hached Hospital & Centre PMI, Sousse | Lower middle income | Paediatric outpatients attending clinic | Children with pharyngitis | 2-8 |
| Seale 2016 | Kenya | Prospective surveillance | Kenya Medical Research Institute, Kilifi County Hospital | Lower middle income | Children admitted for medical care at the hospital | Children located in the area with confirmed iStrep A | 0-12 |
| Tapia 2016 | Mali | Prospective surveillance | 4 Public schools in Djikoronni-para & Sebenikoro | Low income | Children attending 1 of the 4 schools | Children with sore throat | 5-16 |
| Tewodros 2005 | Ethiopia | Prospective surveillance | Black Lion Hospital & 3 elementary schools in Addis Ababa | Low income | Pediatric patients attending the hospital & schools within area | Healthy children at schools & those with confirmed ARF, RHD, APSGN & impetigo | <18 |

iStrep A, invasive Group A Streptococcus; ARF, acute rheumatic fever; APSGN, acute poststreptococcal glomerulonephritis

The country of each article was recorded, with 2 articles obtained from Ethiopia (24, 31), South Africa (14, 32), Tunisia (33, 34) and one article from Kenya (35) and Mali (36). All the studies included in this review made use of the gold-standard, *emm*-typing molecular procedure proposed by Beall (25) and the CDC (26).

### Prevalence of Strep A emm clusters

Five countries within Africa contributed *emm* cluster data to this review (Figure 2). The final dataset included 1,532 isolates representing 126 heterologous *emm* types. Of these, 1,291 isolates, comprising 96 *emm* types, constituted 16 *emm* clusters. Of those remaining, 186 isolates contained 18 single-isolate *emm* clusters (15 *emm* types (143 isolates) representing 15 *emm* clusters belonging to clade Y, while *emm*55, *emm*95 and *emm111* constituted outliers (43 isolates)) (Table 3). The remaining 12 emm types (55 isolates) are amongst those as yet not classified. The predominant clusters were E6 with 294 isolates (18.4%), followed by E3 (n=243, 15.2%) and E4 (n=225, 14.1%). The *emm* clusters with the least number of isolates are D1 and E5, respectively having a single isolate. *emm* Cluster A-C1 was not represented. Sixty-three isolates were reported as ‘untypable’ by authors, thus not assigned an *emm* type, or an ‘old’ *emm* type that does not correspond with the CDC classification.

**Figure 2.**
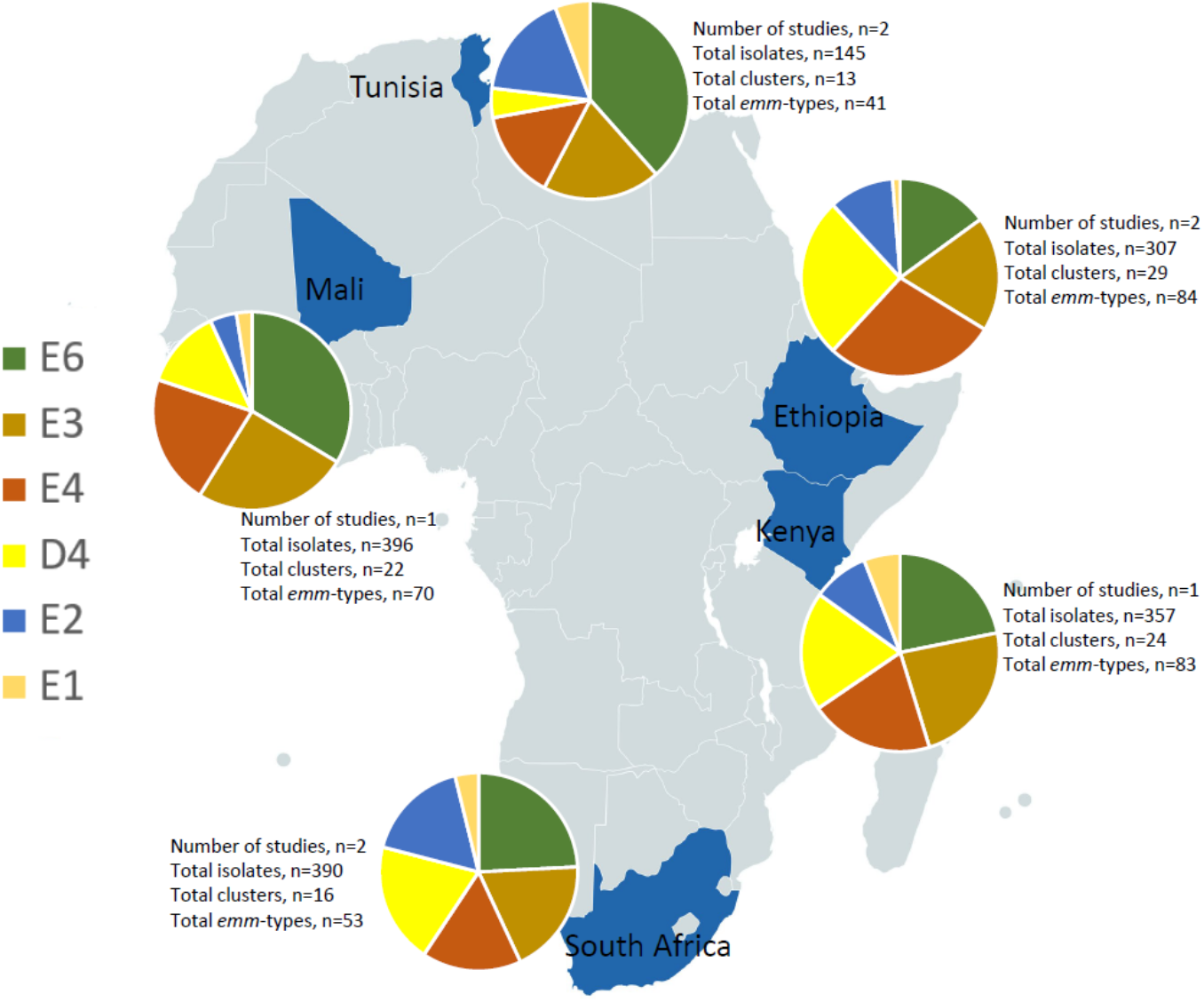
The five countries included in the review, representing the most abundant *emm* clusters.

**Table 3.**
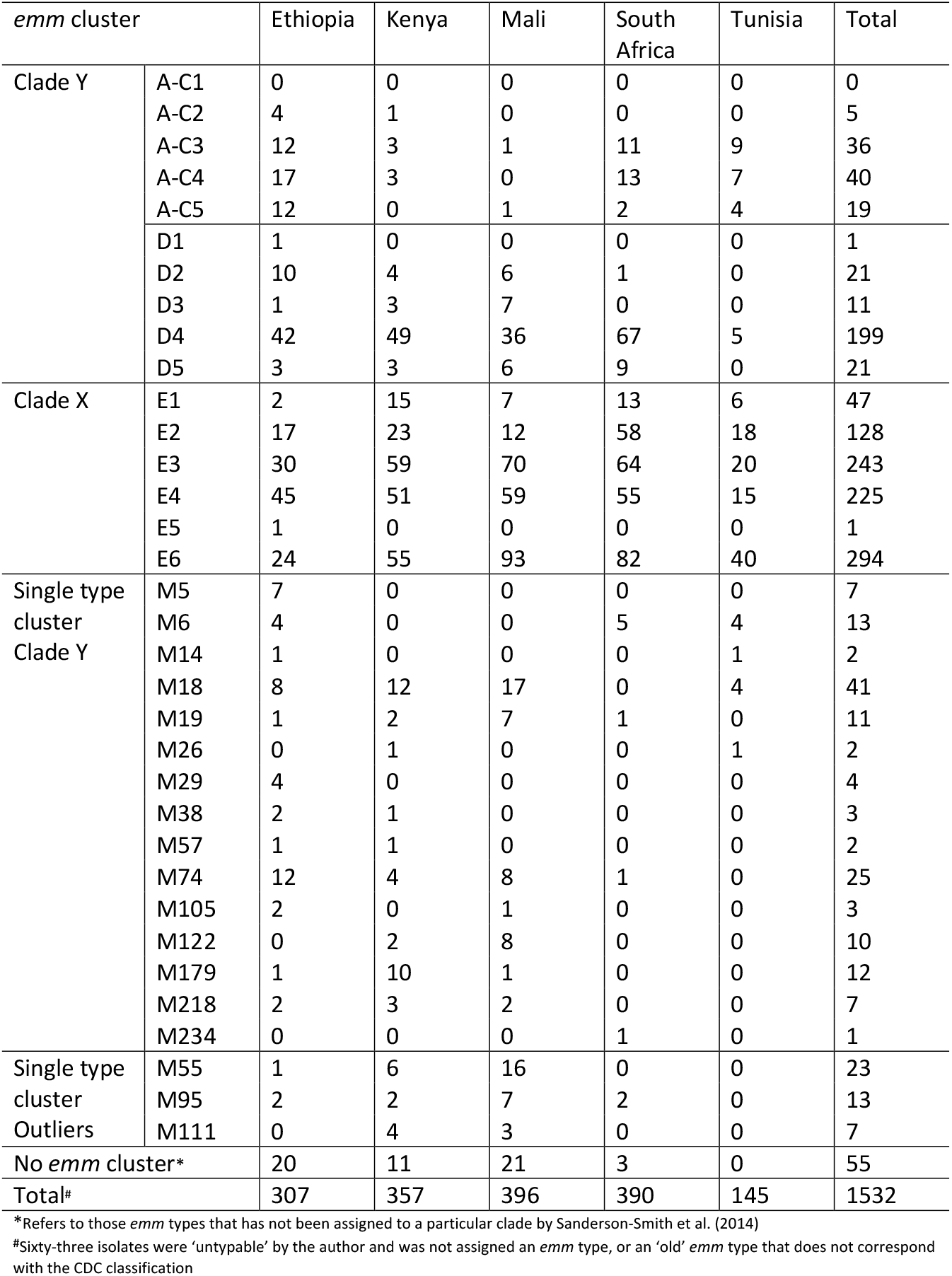
The *emm* cluster distribution, representing the five countries included into the review and their respective isolate count

There were four regions represented across Africa. Variation of clusters across the regions was not remarkable. Interestingly, single-isolate cluster M55 was specific to Mali in West Africa, containing 16 isolates. The highest single-isolate cluster, M18 (n=41 isolates), was not represented in South Africa. Where age of participants in studies was provided, there was no difference amongst children (<18 years of age) in terms of cluster prevalence (Figure 2S-Appendix). By clinical manifestation, isolates from invasive disease numbered 516 (32.4%) (FigureS1-Appendix) with the most prevalent clusters being E3 (n = 91 isolates) followed by E6 (n=82) and D4 (n=77). No variation in *emm* clusters by socio-economic status was apparent.

### Overall Prevalence of Strep A emm clusters represented by the emm types included in the 30-valent vaccine

Cluster E6 was the most represented *emm* cluster (17.97% (95% confidence interval (CI) 12.6% to 24.0%) amongst African isolates (figure 3A). This was followed by E3 [14.17% (95% CI 11.2; 17.4)], E4 [12.6% of isolates (95% CI 9.5; 16.0)], D4 [10.88% (95% CI 6.9; 15.5%)] and E2 [9.12% (95% CI 4.6; 14.9)] of isolates (figure 3B-D). Clusters A-C3, A-C4, A-C5 and E1 each have an effect size of ~2% (Table 4). Isolates from invasive disease were abundant in clusters D4, E2, E3 and E4 while only E6 had a preponderance of strains from non-invasive disease.

**Figure 3.**
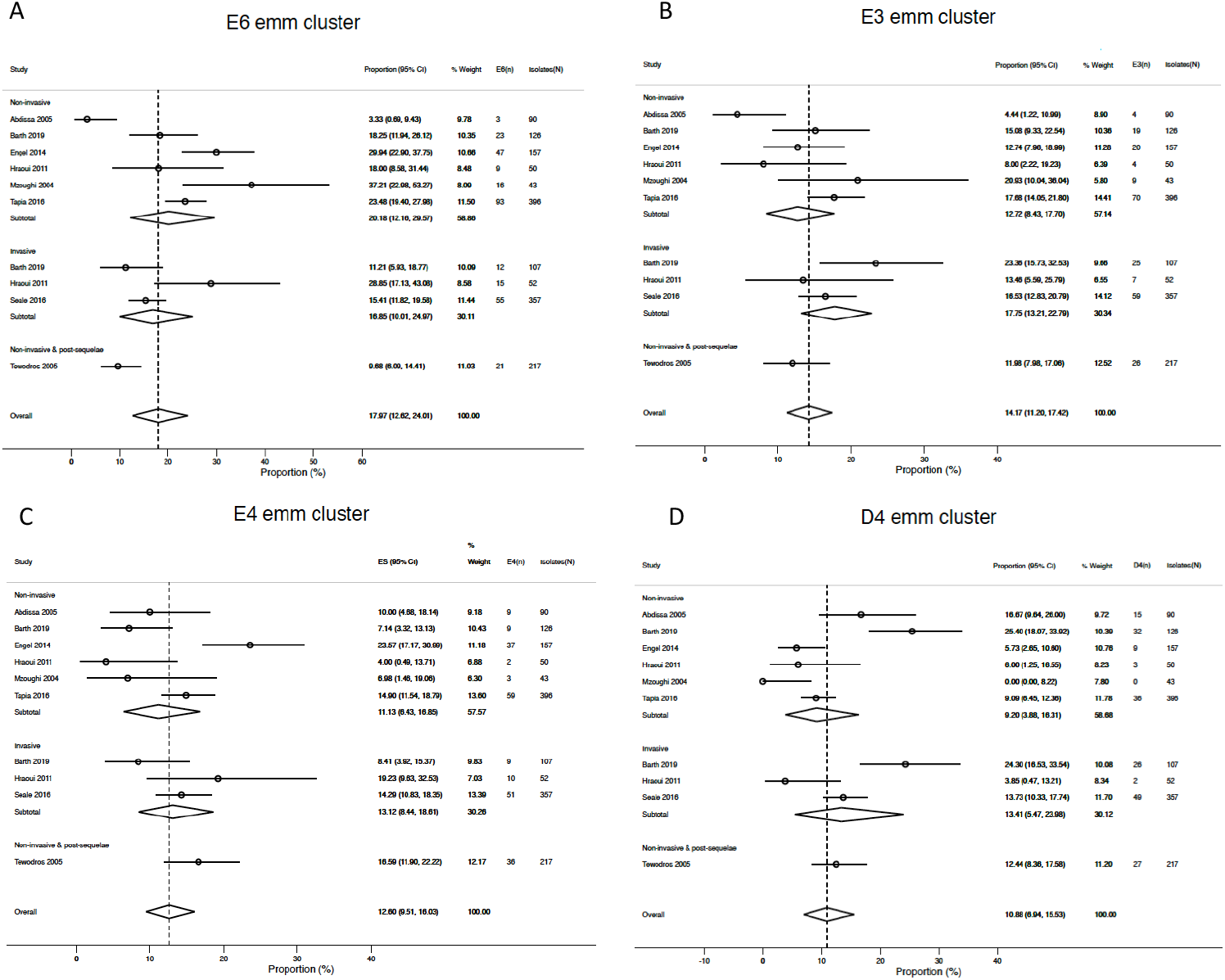
Forest plots showing the combined prevalence estimates of the four most abundant *emm* clusters of all included studies. A, E6; B, E3; C, E4; D, D4

**Table 4.** Summary of the meta-analyses completed^#^

| Cluster | *emm* types in vaccine | *emm* types in African isolates | Combined prevalence (95% CI) | Number of *emm* type isolates |  |  |
| --- | --- | --- | --- | --- | --- | --- |
|  |  |  |  | NINV (%) | INV (%) | Total |
| E6 | 11, 75, 81 | Prot: 11, **48, 59, 65**, 75, 81, **85, 94** (*n*=235 (79.9%));<br>N/P: 42, 63, 67, 99, 158, 182;<br>NINV: *n*=160 (83.8%); INV: *n*=59 (72.0%)) | 18.0% (12.62; 24.01) | 191 (65.0) | 82 (27.9) | 294 |
| E3 | 44, 49, 58, 87, 118 | Prot: **9, 15, 44**, 49, 58, **79, 87, 118** (*n*=130 (53.5%));<br>N/P: 25, 61, 82, 103, 113, 180, 183, 209;<br>NINV: *n*=73 (57.9%); INV: *n*=47 (51.6%)) | 14.2% (11.20; 17.42) | 126 (51.9) | 91 (37.4) | 243 |
| E4 | 2, 22, 28, 73, 77, 89, 114 | Prot: 2, **8, 22, 28, 73, 77, 89, 109, 114** (*n*=202 (89.8%));<br>N/P: 84, 112, 124, 175, 225;<br>NINV: *n*=115 (96.6%); INV: *n*=52 (74.3%)) | 12.6% (9.51; 16.03) | 119 (52.9) | 70 (31.1) | 225 |
| D4 | 83 | Prot: **33, 83** (*n*=15 (7.5%));<br>N/P: 41, 43, 53, 56, 64, 70, 80, 86, 93, 98, 116, 119, 121, 186, 192, 208, 223, 224, 230;<br>NINV: *n*=5 (5.3%); INV: *n*=9 (11.7%)) | 10.9% (6.94; 15.53) | 95 (47.7) | 77 (38.7) | 199 |
| E2 | 92 | Prot: **66, 68, 76, 92, 102** (*n*=92 (71.9%));<br>N/P: 50, 90, 104, 106, 110, 168;<br>NINV: *n*=49 (77.8%); INV: *n*=38 (69.1%)) | 9.1% (4.61; 14.86) | 63 (49.2) | 55 (43.0) | 128 |
| E1 | 4, 78 | Prot: 4, 78 (*n*=31 (66.0%));<br>N/P: 60, 165;<br>NINV: *n*=22 (78.6%); INV: *n*=8 (47.1%)) | 1.9% (0.62; 3.81) | 28 (59.6) | 17 (36.2) | 47 |
| A-C4 | 12 | Prot: 12 (*n*=35 (87.5%)); I (*n*=3 (42.9%));<br>N/P: 39, 193, 229;<br>NINV: *n*=17 (94.4%); INV: *n*=3 (42.9%)) | 2.1% (0.37; 4.81) | 18 (45.0) | 7 (17.5) | 40 |
| A-C3 | 1 | Prot: 1 (*n*=32 (88.9%));<br>N/P: 238;<br>NINV: *n*=21 (95.5%); INV: *n*=1 (25.0%)) | 2.0% (0.46; 4.31) | 22 (61.1) | 4 (11.1) | 36 |
| A-C5 | 3 | Prot: 3 (*n*=19 (100%));<br>N/P: NA;<br>NINV: *n*=17 (100%); INV: *n*=2 (100%)) | 0.9% (0.01; 2.72) | 17 (89.5) | 2 (10.5) | 19 |

<sup>#</sup>301 isolates (19.6%) comprising 39 *emm* types are not included in any of the *emm* clusters contained in the 30-valent vaccine; these include 60 isolates representing seven *emm* clusters; 186 isolates representing single-isolate clusters & 55 isolates were not classified as according to Sanderson-Smith. Bold *emm* types represent cross-opsonized non-vaccine types. Study completed by Tewodros & Kronvall (2005) did not clearly differentiate its *emm* types according to clinical manifestation. Combined prevalence calculated with M-H meta-analysis procedures. NINV, non-invasive; INV, invasive; Prot, protected; N/P, not protected; NA, none.

### M Protein Vaccine Coverage

Just over eighty percent (80.3%) of African Strep A isolates are classified in clusters included in the 30-valent vaccine (Figure 4). However, based on *emm* types within the vaccine, together with *emm* types known to be cross-opsonized, the number of African Strep A isolates that potentially could be covered by the 30-valent vaccine amounts to 892 of the 1532 isolates corresponding to 58.22% (comprising 599 vaccine type *emm* types and 293 non-vaccine *emm* types) (11). For the *emm* types representing the remaining 640 isolates (41.78%), there is either no information yet available about possible cross-protection, or the *emm* types would not be expected to cross-react with the 30-valent vaccine antisera because they are in single-*emm* clusters or in clusters not represented by the vaccine. Interestingly, isolates classified as *emm*30 (AC-2), *emm*36 (D1), *emm*51 (E5) and *emm*97 (D5), despite not being in a cluster represented in the vaccine, are nevertheless afforded cross-protection.

**Figure 4.**
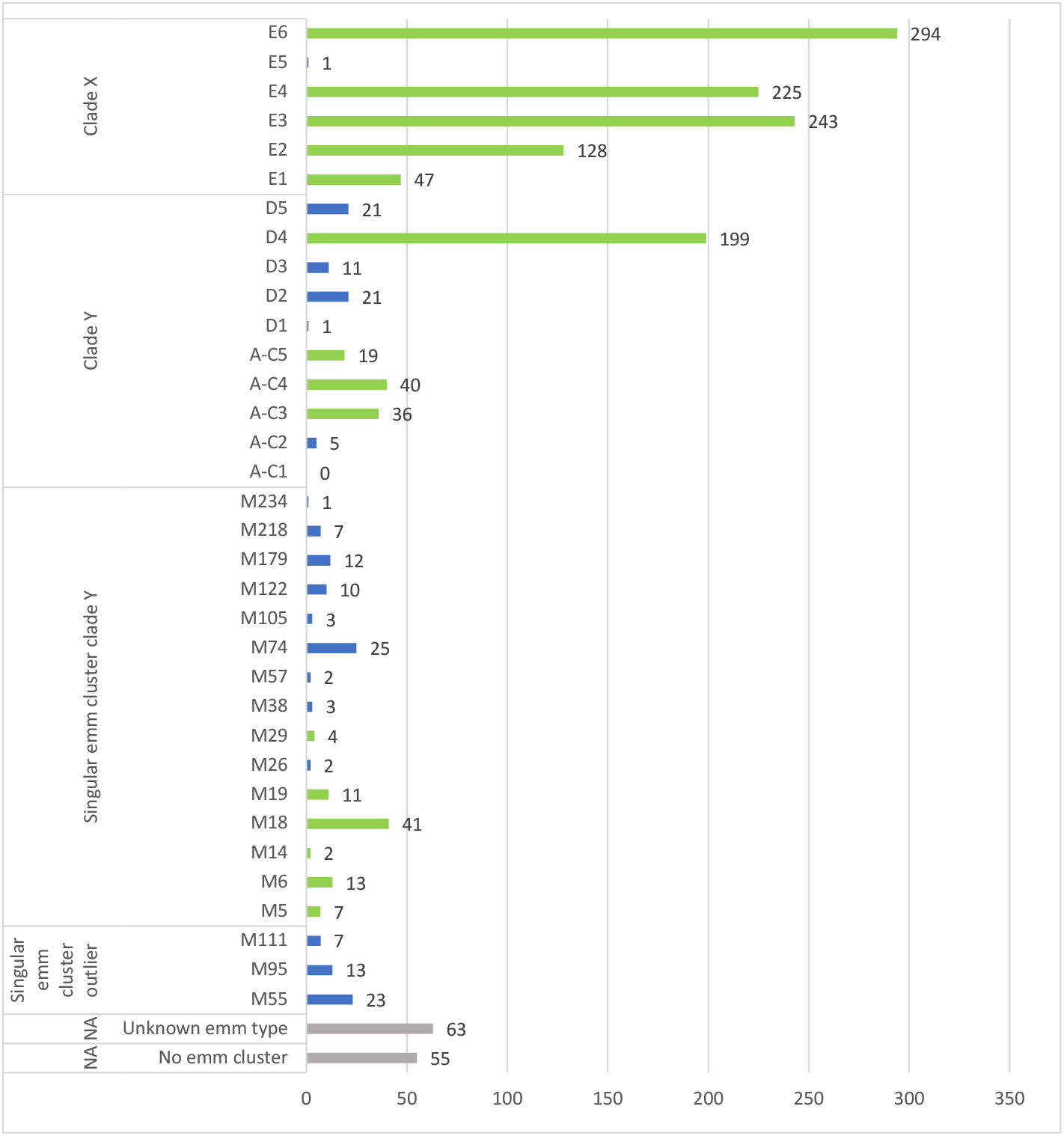
Count of *emm* clusters from African studies included in this systematic review. Green bars indicate *emm* clusters represented in the 30-valent vaccine. Blue bars represent *emm* clusters not included in the vaccine. Grey bars represent isolates unassigned to a cluster or were ‘untypable’ according to the authors’ report. Numbers represent the count of isolates across all studies.

With regard to invasive *emm* types in Africa, the overall potential coverage of the vaccine (based on published results of cross-opsonization) was 54.1% for clusters included in the meta-analyses (Table 4). More specifically, coverage for clusters E6, E4 and E2 ranges from 69-74% of invasive isolates; only ~50% of strains would be protected in E3 and coverage for the remaining clusters were below 47% except A-C5 (100% coverage) as there were only two invasive strains reported. Interestingly, the 30-valent vaccine would potentially only provide 12% coverage to invasive isolates belonging to the fourth highest cluster, D4 (n=28 *emm* types).

### Assessment of risk of bias of included studies

The results from the assessment is portrayed in the Table 5, with two studies having a low risk of bias (32, 36) and the remaining studies being of moderate bias. All the articles narrowed down their target population by focusing on a specific age group, clinical manifestation or geographical area. The data in all included studies were collected directly from the study participants as opposed to by proxy, confirming the reliability of sample collection and patient demographics. The included studies clearly described the phenotypes of patients, providing an acceptable case definition or diagnostic algorithm. Studies focusing on invasive Strep A infections isolated Strep A from normally sterile sites such as blood, cerebrospinal fluid, joints, bones or synovium amongst others. Non-invasive Strep A was isolated from skin or throat via swabs of the infected area.

**Table 5.**
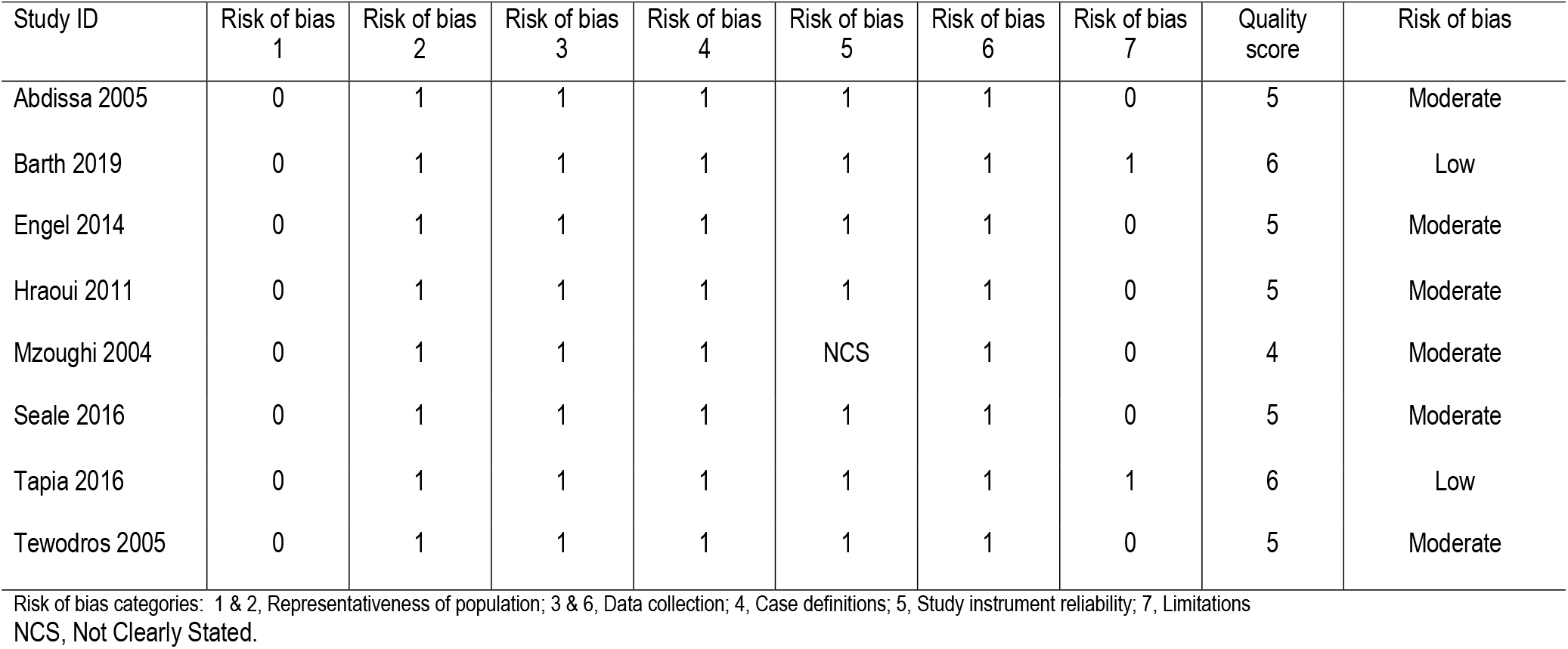
Summary of risk of bias assessment

## Discussion

This systematic review provides evidence for the distribution of *emm* clusters of Strep A in Africa, specifically focusing on the epidemiological differences within Africa and added value of the *emm* clustering system in assisting with vaccine development. Using prevalence data obtained from eight studies representing five countries within Africa, this report identified the predominant *emm* clusters in Africa, namely E6 followed by E3, E4 and D4. We further report that the *emm* clusters contained in the current 30-valent vaccine could provide considerable coverage across the diversity of *emm* cluster types in Africa.

Comparing results to other *emm* clustering epidemiology studies, it is clear that there are variances amongst the dominant *emm* clusters between regions. Only cluster E3 in the present study is common with the Pacific region (17). Within the USA, E4 is the third highest cluster, whereas A-C3 and A-C4 together only amount to ~2% of the total strains isolated in Africa (16). This study emphasizes that *emm* clusters E6, E3 and D4, prevalent in the African populations where the burden of Strep A infections is highest (37), should take prominence alongside clusters E4, A-C3 and A-C4. We note that there are a number of *emm* clusters containing a single *emm* type as they do not share similar binding properties or sequences. Also, there are many *emm* types that have as yet not been categorized into a particular cluster, as this may be due to their recent emergence post the proposed cluster system. This should be the focus of future studies in which more associations with human host protein binding could be tested to determine any other similarities between single-*emm* clusters.

Steer reported that the African and Asian regions had the greatest diversity of *emm* types (13). This could be due to a variety of factors causing site-tissue tropism and disease manifestation, promoting the dominance of heterologous *emm* types in different regions (38). Our review provides no evidence for marked variation across the continent amongst most of the more prominent *emm* clusters. When considering the ages of participants infected with Strep A, there appears to be no differences compared to that of the overall estimates. There is an increased risk for the transmission of Strep A in poorer countries due to household crowding and the lack of income for proper healthcare (39). Evaluating socio-economic status amongst our studies revealed little to no differences in *emm* cluster data.

Amongst non-invasive infections, cluster E6 was the most abundant cluster. This is in accordance with previous reports completed by Sagar (40), Dhanda (41) and Arêas (42), that identified *emm* types belonging to cluster E6 (emm75, emm81) as the predominant isolates obtained from countries closely relating to the impoverished environments within Africa, India and Brazil respectively. However, when referring to invasive disease, the predominant *emm* clusters are E3, followed by E6 and D4, which is complimentary to the *emm* cluster data shown by Chiang-Ni (18).

In terms of the current 30-valent vaccine (11), with the assumption that the *emm* type prevalence data from the eight included studies could be generalised for the entire continent, vaccine coverage would be 55.92% of strains isolated in Africa. Frost had shown cross-reactive protection of a single *emm* type with the remaining *emm* types within the same cluster, specifically that of E4 (19). Thus, hypothetically assuming that if a single *emm* type in the 30-valent vaccine would provide cross-protection to the remaining isolates within the cluster, a *emm* cluster-based vaccine would then extend coverage to ~80% protection against Strep A (Figure 4). Of interest cluster D4, which comprises 28 heterologous *emm* types, and ranked high in this analysis, has only a single representation (*emm*83) included in the vaccine. If cross-protection were to occur within clusters, more *emm* types belonging to cluster D4 ought to gain a particular importance for inclusion into new vaccines, especially since D4 (10.9% of isolates) is the fourth highest abundant cluster within Africa. It is also important to note that coverage extended to invasive isolates was sub-optimal (n=219, 54.1%).

One of the main strengths of this review is attributed to the use of multiple databases searched, using an African search filter and a robust approach to the meta-analysis of the data. We systematically and purposefully assessed all the data available with no language exclusions, or restrictions to a clinical manifestation of disease, using the most recently published standard quality assessment tools for prevalence studies. We also assessed the risk of bias present in the individual articles, showing that the quality was reasonably high, thus allowing for comparisons across the studies. The main limitations of the review are due to the lack of epidemiological data obtained from low to middle income countries in Africa, especially given their relatively high burden of Strep A infections. The inclusion of more articles reporting on the prevalence of Strep A may further assist in distinguishing differences amongst the geographical location, age and socio-economic categories. A further limitation to the results of our systematic review is the significant heterogeneity in the prevalence estimates produced in the meta-analysis, however, this is expected when pooling prevalence studies. We made use of the Freeman-Tukey double arc-sine transformation to stabilize the variance of primary studies before pooling, thus limiting the impact of studies with either small or large prevalence on the overall pooled estimates, as well as across major subgroups (30).

## Conclusion

In conclusion, this systematic review provides the latest evidence for the distribution of *emm* clusters of Strep A in Africa. We show that there is negligible variation in emm clusters as regards regions, age and socio-economic status across the continent. We further report that the current 30-valent vaccine will provide considerable coverage across the diversity of *emm* cluster types in Africa, thus providing direction for future work to include coverage of clusters D4, E2-E4 and E6, given that they comprise 83% of the total isolates obtained in Africa.

## Acknowledgements

This work is based on the research supported in part by the National Research Foundation of South Africa (Grant Numbers: 116287). Any opinion, finding, conclusion or recommendation expressed in this material is that of the author(s) and the NRF does not accept any liability in this regard. MEE, TS, KE and BM are supported by a grant, NW17SFRN33630027 from the American Heart Association, United States.

MEE, TS and AM wrote the protocol designed the study. TS implemented and managed the study. TS and KE reviewed articles and recorded the data. BM was the third arbitrator. TS and MEE conducted statistical analysis. TS, JBD and MEE interpreted the data and wrote the manuscript. All authors read and approved the final manuscript.

## Appendix – Tables and Figures

**Table S1.**
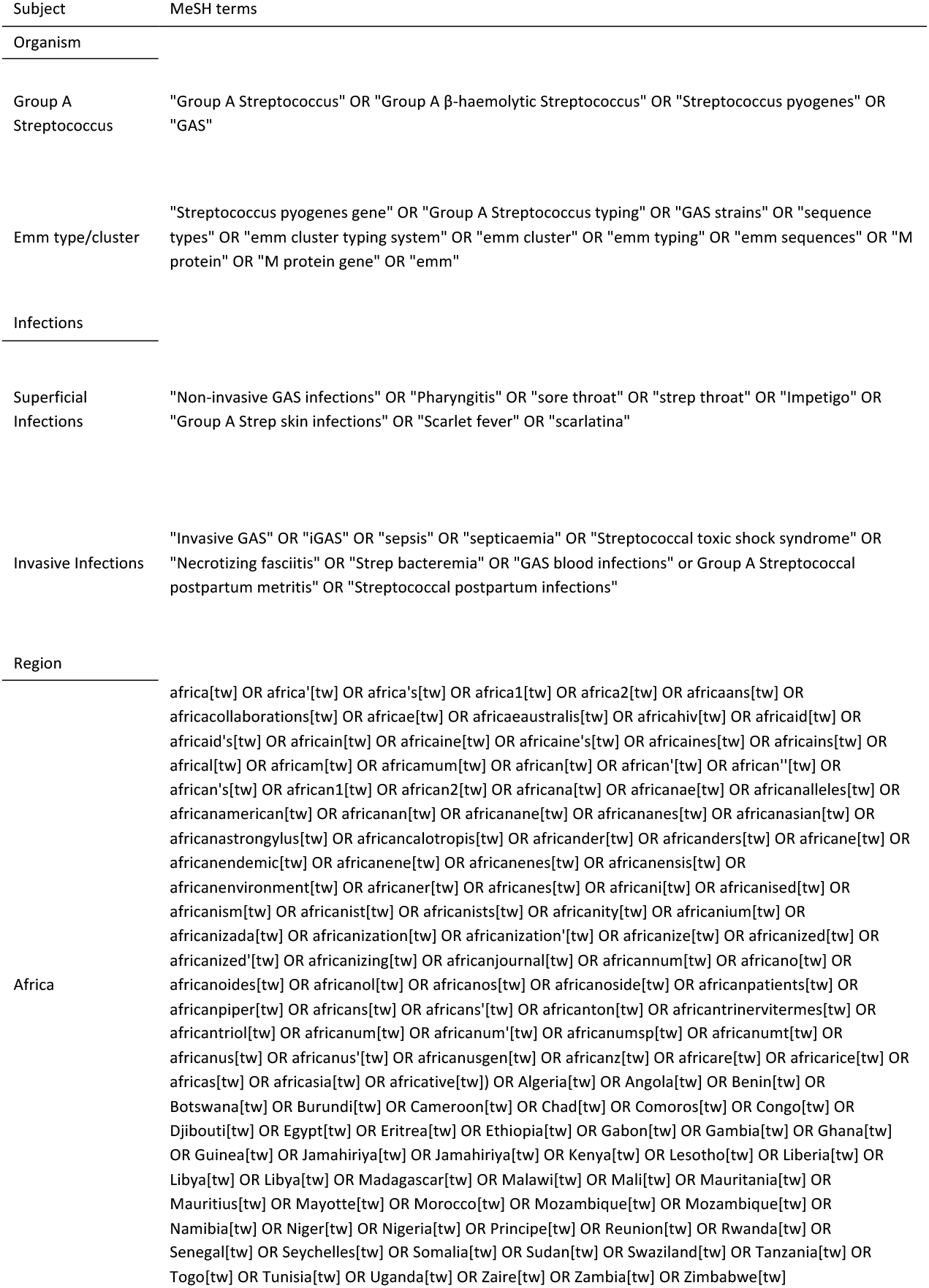
Search Strategy with MeSH terms used for databases

**Table S2.** Inclusion criteria

| Inclusion Criteria |  |
| --- | --- |
| Study aims | Studies that describes the distribution of *emm*-types or -clusters in a population |
| Detection method | All study designs in which a Strep A *emm*-typing scheme was performed, PCR amplification of the *emm* gene or whole genome sequencing |
| Participants | All participants with confirmed Strep A infection at the time of enrolment |
| Region | Countries situated within the African continent |
| Publication types | Published and unpublished studies |
| Language | All, with full English abstracts |

**Table S3.**
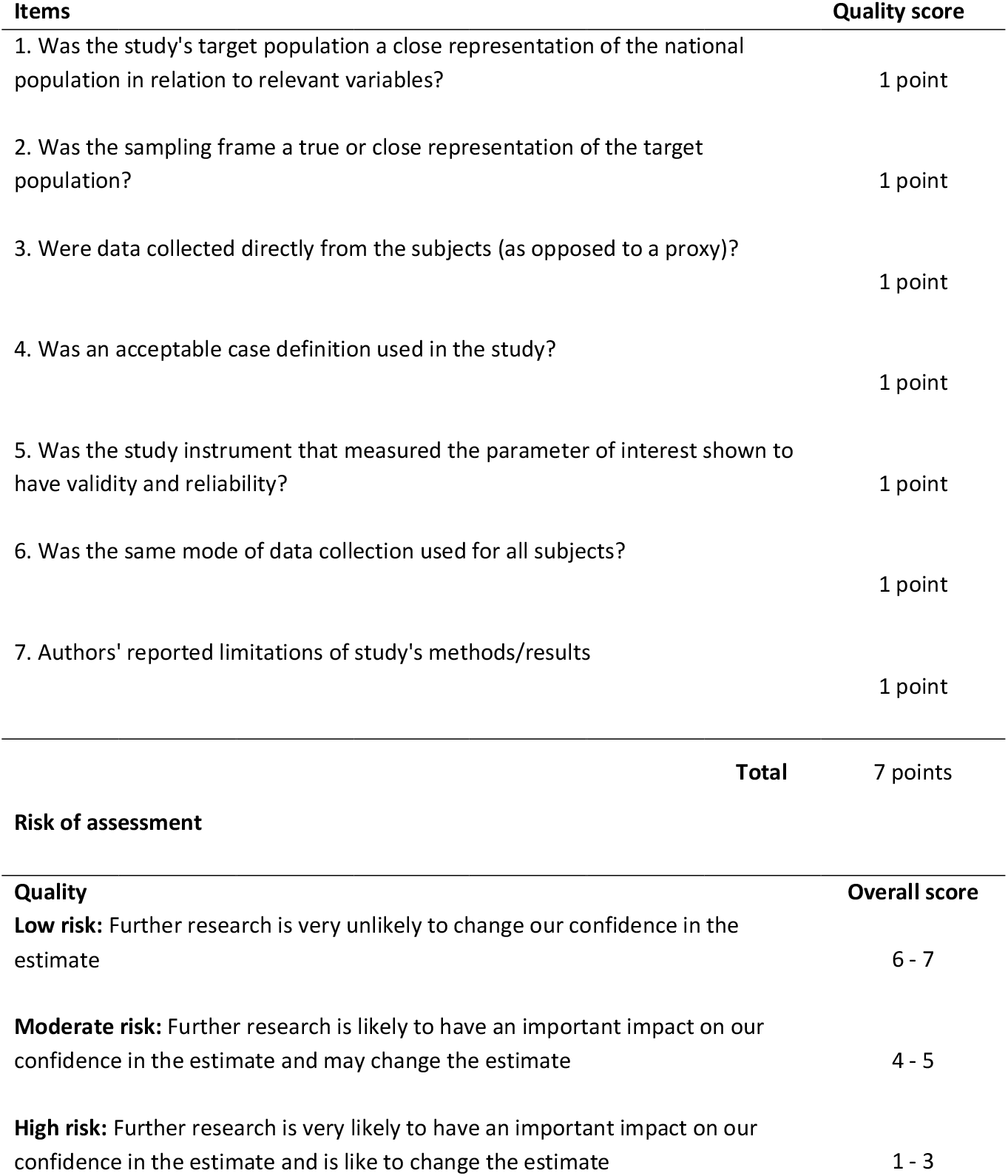
Study appraisal for risk of bias for included studies adapted from Hoy (28) and Werfalli (22).

**Table S4.**
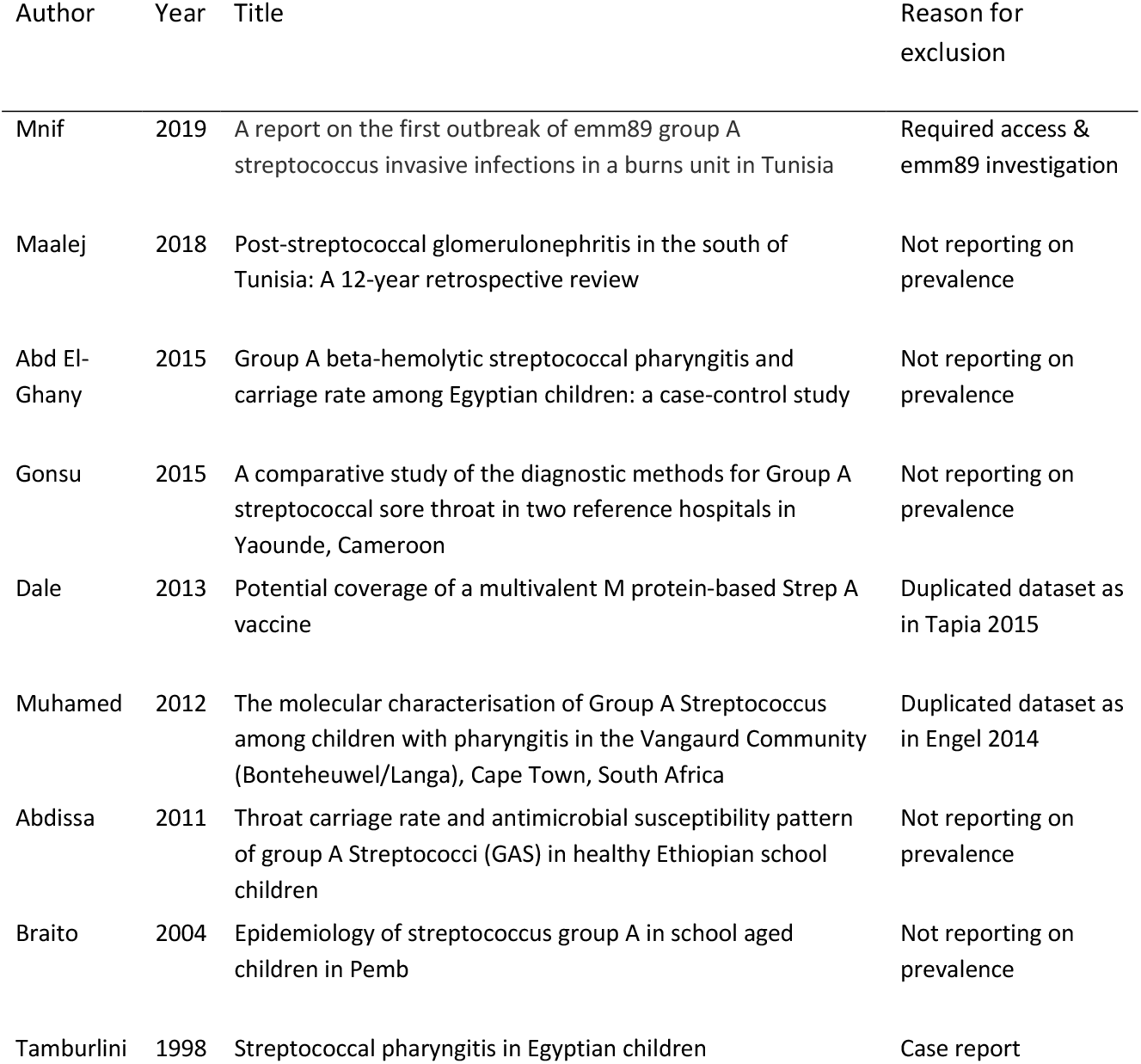
Characteristics of excluded studies

**Figure S1.**
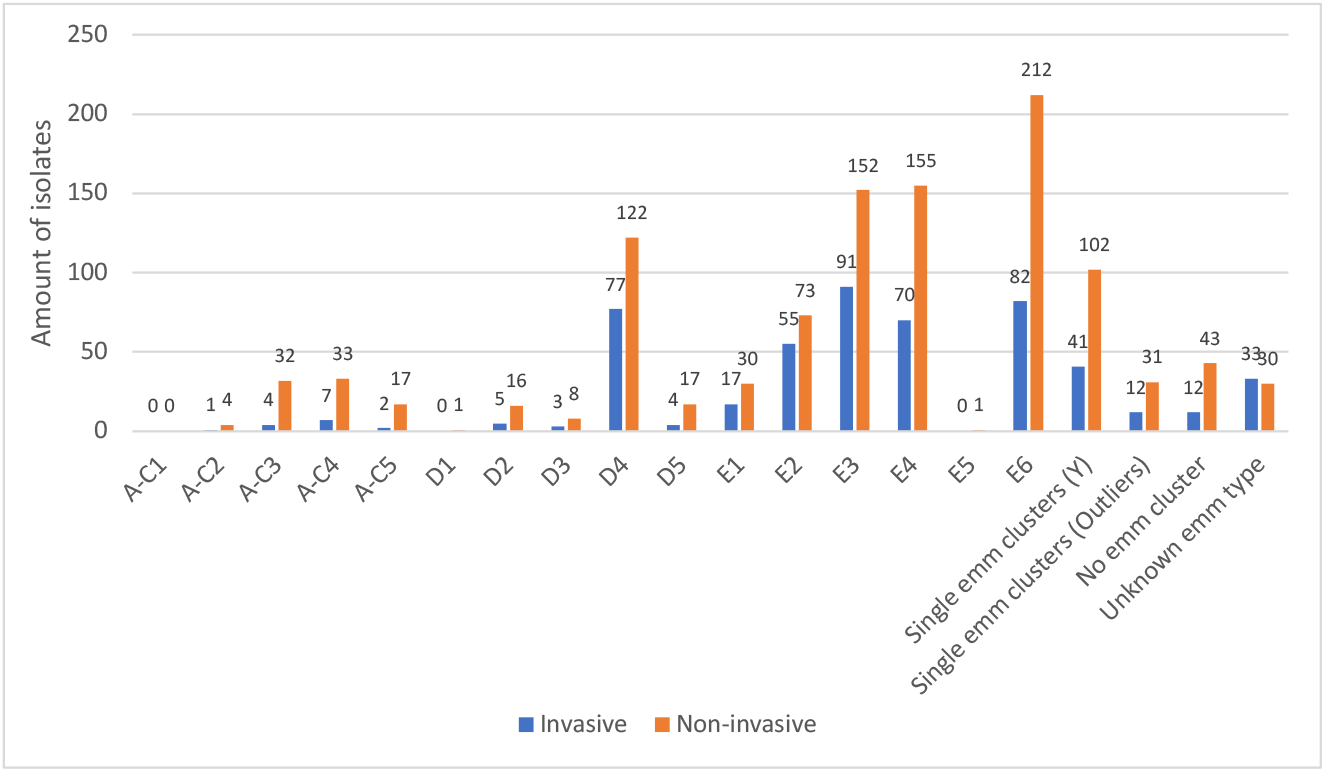
Total prevalence of *emm* clusters categorized based on clinical manifestation. Invasive isolates shown in blue (n=3 studies) and non-invasive isolates shown in orange (n=6).

**Figure S2.**
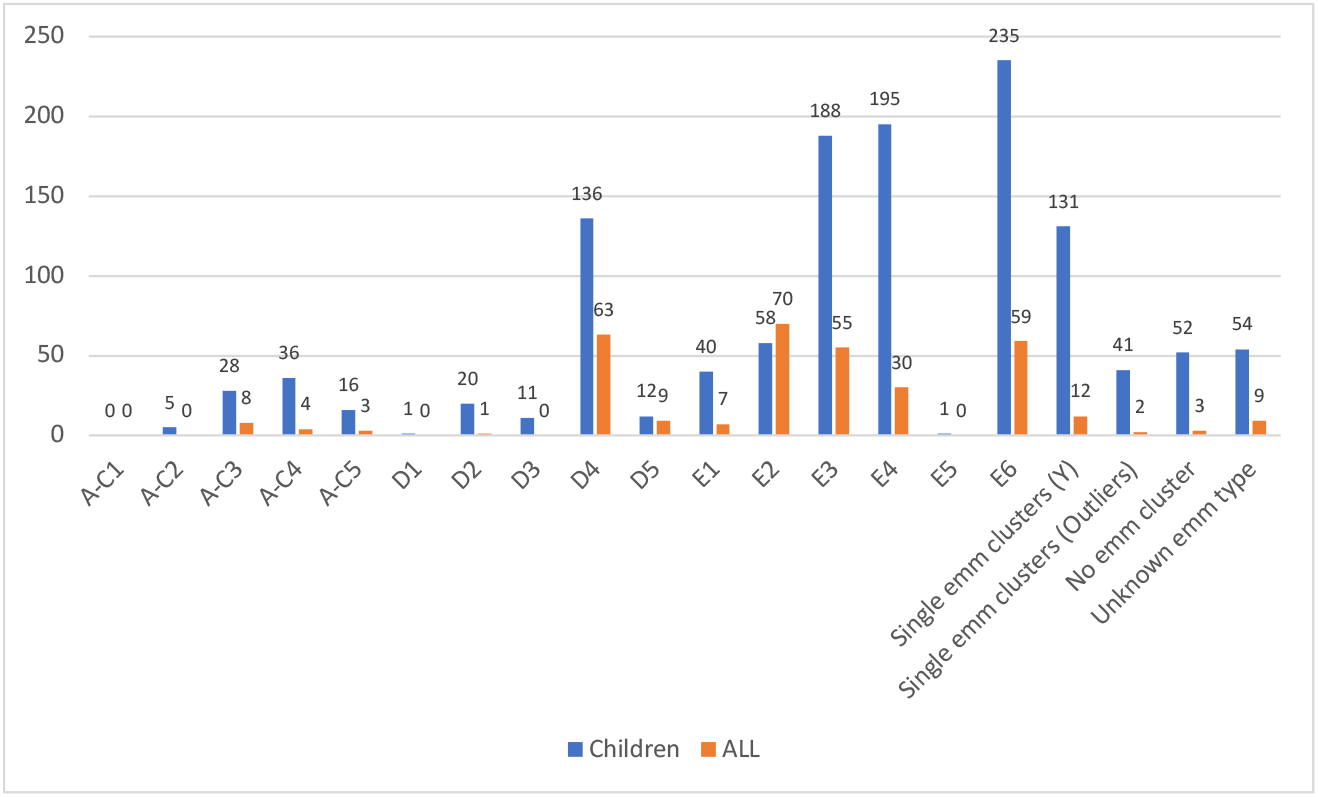
Total prevalence of *emm* clusters categorized based on age of participants included in each study. Children (<18 years of age) shown in blue (n=7 studies) and participants of all ages shown in orange (n=2).

